# 3,4-hydroxyphenyl lactic acid from *Lactiplantibacillus plantarum* prevents alcoholic liver injury in mice

**DOI:** 10.1101/2025.03.12.642873

**Authors:** Caixia Zhang, Yirong Chen, Shijia Huang, Xiaofeng Bian, Bowen Yang, Siyan Lu, Xueting Fu, Wei Zhao, Yan Pan, Shuli Zhao

## Abstract

Alcoholic liver disease (ALD) is a major health burden linked to oxidative stress, gut dysbiosis, and disrupted hepatic metabolism. While *Lactiplantibacillus plantarum* (*L. plantarum*) has shown potential in alleviating ALD, the specific mechanisms and bioactive components remain unclear. This study investigated the hepatoprotective effects of *L. plantarum* fermentation liquid (PFL) and its key metabolite, 3,4-hydroxyphenyl lactic acid (HPLA), against alcohol-induced liver injury. Using acute and chronic alcohol intoxication mouse models, we demonstrated that PFL significantly reduced mortality, attenuated hepatocyte damage, and restored alcohol dehydrogenase (ADH) and aldehyde dehydrogenase (ALDH) activities. UHPLC-QTOF-MS/MS analysis identified HPLA as the primary active component in PFL, exhibiting potent antioxidant properties. In vitro and in vivo experiments revealed that HPLA mitigated oxidative stress by enhancing superoxide dismutase (SOD) and glutathione peroxidase (GSH-Px) activities, reducing malondialdehyde (MDA) levels, and suppressing lipid accumulation. Mechanistically, network pharmacology and molecular validation highlighted that HPLA alleviated hepatic injury by modulating the EGFR/PPAR-α signaling axis, thereby counteracting alcohol-induced oxidative stress and lipid metabolism disorders. These findings elucidate a novel “gut-liver” axis mechanism mediated by HPLA, offering a theoretical foundation for the clinical application of *L. plantarum* and its metabolites in ALD management.

**Importance:** Numerous animal studies and clinical trials have demonstrated the effectiveness of probiotics in treating alcoholic liver disease. In this study, we primarily created a mouse model for both acute and chronic alcoholism and discovered that *Lactiplantibacillus plantarum* fermentation solution significantly decreased the inflammatory response and oxidative stress caused by alcohol. Its key metabolite, 3,4-hydroxyphenyl lactic acid (HPLA), exhibited strong antioxidant properties, helping to reduce oxidative stress by preventing lipid accumulation. This research offers insights into how probiotic interventions can mitigate alcoholic liver damage, enhancing our understanding of the protective effects of *Lactiplantibacillus plantarum* and emphasizing the potential of its metabolite, HPLA, as a targeted treatment for alcoholic liver disease.

## Introduction

Excessive alcohol consumption poses serious health risks, leading to a range of acute and chronic conditions, including liver disease, gastrointestinal disorders, and gut microbiome disorders (Shukla et al. 2018; Xie et al. 2021; Simon et al. 2022). Alcohol metabolism produces reactive oxygen species that cause oxidative stress and inflammation, leading to cell damage and organ dysfunction (Shi et al. 2015; Chen et al. 2024). In addition, alcohol disrupts the integrity of the intestinal barrier, leading to increased permeability, harmful substances entering the bloodstream, and the exacerbation of inflammation throughout the body (Martino et al. 2022). Studies have also suggested that long-term alcohol intake can lead to an imbalance in gut flora, further aggravating liver damage. This ‘gut-liver’ axis plays a key role in the onset, progression, and worsening of alcoholic liver disease (Li et al. 2020).

*Lactiplantibacillus plantarum* (*L. plantarum*) is a lactic acid bacterium that predominantly colonizes the gut and positively affects the intestinal microbial environment. Currently, *L. plantarum* is widely used in food production, industry, and healthcare, and it has great potential for further research and development (Gam et al. 2022). Prolonged alcohol consumption, which can deplete the intestinal flora, particularly reduces the population of *L. plantarum* (Kirpich et al. 2008). Yet studies have suggested that this bacterium has protective effects against alcoholic liver injury. The underlying molecular mechanisms, however, remain unknown.

Privous study showed that *L. plantarum* could activate the Keap-Nrf2-ARE signal pathway by upregulating the Nrf2 expression to protecte the organism from oxidative stress and maximally reduce liver injury (Liu et al. 2019). *L. plantarum* ST-III culture supernatant can alleviates acute alcohol-induced liver damage by inhibiting oxidative stress and endoplasmic reticulum stress via one mechanism (Zhou et al. 2020), but, the specific molecules involved in antioxidant stress are not clear.

This study involved the development of animal models of both acute and chronic alcohol intoxication to analyze the protective effects of *L. plantarum* against alcohol toxicity. Ultra-high-performance liquid chromatography-quadrupole-time of flight mass spectrometry (UHPLC-QTOF-MS/MS) was employed to identify the key active components of *L. plantarum* fermentation liquid (PFL). The protective mechanism of the identified component, 3,4-hydroxyphenyl lactic acid (HPLA), was then validated using both in vivo and in vitro models of alcohol intoxication. This study provides a solid theoretical foundation for the protective effects of *L. plantarum* in alcohol-induced liver injury via ‘gut-liver’ axis, and experimentally validates the in vivo functions and potential mechanisms of the active ingredient HPLA of *L. plantarum*. Furthermore, we explore strategies to optimize its application, thereby maximizing its potential health benefits.

## Materials and Methods

### L. plantarum Preparation

*Lactiplantibacillus plantarum* (ATCC 8014) was purchased from the China General Microbiological Culture Collection Center (CGMCC), Beijing, China. These bacteria were cultured in a specific liquid medium composed of 1% honey, 4% brown sugar, and water, at 37°C. When the bacterial concentration reached 107, the bacteria and culture supernatants were collected for centrifugation experiments.

### Animal Models

Eighty environmentally acclimatized 6-week-old Kunming mice (40 males and 40 females), each weighing approximately 30 g, were obtained from the Experimental Animal Center (Kalaisi). Mice were randomly assigned to four groups: control, alcohol (model), *L. plantarum* (P), and *L. plantarum* fermentation liquid (PFL). For the mouse survival analysis model, there were eight mice in each group, totaling 32 mice, and for the mouse acute alcohol intoxication model, there were six mice in each group, totaling 24 mice. Similarly, in the chronic alcoholic liver injury model, six mice were used in each group, for a total of 24 mice.

### Righting reflex

Six mice from each group were used for behavioral observation. Intoxication was assessed based on the loss of the righting reflex(Chen et al. 2014). After alcohol administration by gavage, mice were placed on their backs every 5 min. If a mouse remained in the supine position for more than 30 s, it was considered to have lost its righting reflex and therefore intoxicated.

### Biochemical analysis

Whole blood was collected from the mice, and the serum was separated. Livers were excised and weighed. A 10% (w/w) liver tissue homogenate was prepared and the supernatant was used for various biochemical assays. Serum levels of aspartate aminotransferase (AST), alanine aminotransferase (ALT), high-density lipoprotein cholesterol (HDL-C), low-density lipoprotein cholesterol (LDL-C), triglycerides (TG), and total cholesterol (T-CHO) were measured using a fully automated biochemical analyzer. The liver activities of acetaldehyde dehydrogenase (ALDH), ethanol dehydrogenase (ADH), malondialdehyde (MDA), superoxide dismutase (SOD), and glutathione peroxidase (GSH-Px) were determined using assay kits from Nanjing Jiancheng Bioengineering Institute, following the manufacturer’s instructions.

### Reverse transcription-quantitative PCR (RT-PCR) analysis

Total RNA was extracted from cultured cells and liver tissues using Trizol reagent (Invitrogen, Waltham, MA), and cDNA was synthesized using PrimeScript ® RT Master Mix (Vazyme, Nanjing, China). Real-time PCR was performed in triplicate on a Biological Systems Q5 system using a SYBR Green PCR Kit (Vazyme). Relative mRNA expression levels were analyzed using the 2^−ΔΔCt^ method. A list of primers used for qRT-PCR is provided in Table S1.

### Liver histological analysis

Fresh liver tissue was fixed in 4% paraformaldehyde at room temperature (20-26℃), dehydrated, and embedded in paraffin. The tissue was then sectioned with a thickness of 4-5 μm and stained with hematoxylin and eosin. Histomorphological evaluation of sections from each group was conducted using a standard light microscope (Olympus BX53; Tokyo, Japan) at 40× magnification.

### UHPLC-QTOF-MS/MS analysis

#### 1. Bacterial liquor alcohol extract

The bacterial solution was centrifuged at 13,000 rpm for 15 min and the supernatant was passed through a 0.22-μm filter membrane. Next, 100 μl of methanol was added and 300 μl of methanol was ultrasounded for 15 min after vortex. Centrifuge at 13,000 rpm for 15 min. Finally, 360 μl of the supernatant was blown dry, 100 μl of 50% acetonitrile was added to redissolve, after which it was centrifuged at 13,000 rpm for 15 min and the supernatant was taken into the sample.

#### 2. Lyophilized dansylhydrazine derivatization

Dansylhydrazine (DnsHz) was dissolved in acetonitrile to obtain a 10 mg/ml solution, while water was used with MES to form a buffer solution (pH 3.5, 0.1 M). EDC and HoAt were prepared into 100 and 10 mM solutions with MES buffer, respectively. The quencher CuCl_2_ was prepared as a 100 mM solution with H_2_O. After freeze-drying, 50 mg of bacterial solution was taken, 1 ml of 50% acetonitrile was added to redissolve, it was centrifuged at 13,000 rpm for 15 min, and then the supernatant was passed through a 0.22-μm filter membrane. Next, 20 μl was added into a 1.5 ml EP tube, 20 μl EDC and 20 μl HOAT were added, and it was blown and mixed well. DnsHz (20 μl) was added and swirled well, and it reacted in a 20°C water bath for 90 min, after which 20 μl CuCl2 solution was added, and it reacted in a 40°C water bath for 30 min, and was then quenched. After the nitrogen was blown dry, 100 μl of 50% acetonitrile was added to redissolve, and the supernatant was taken into the sample after being centrifuged at 13,000 rpm for 15 min.

#### 3. Bacterial solution direct dansylhydrazide derivatization

The bacterial solution was centrifuged at 13,000 rpm for 15 min and the supernatant was passed through a0.22-μm filter membrane. Then 20 ml was added into a 1.5 ml EP tube, 20 μl EDC and 20 μl HOAT were added, and it was blown and mixed well. DnsHz (20 μl) was added and swirled well, and it reacted in a 20°C water bath for 90 min, after which 20 μl CuCl_2_ solution was added, and it reacted in a 40°C water bath for 30 min, and was then quenched. After the nitrogen was blown dry, 100 μl of 50% acetonitrile was added to redissolve, and the supernatant was taken into the sample after being centrifuged at 13,000 rpm for 15 min.

#### 4. Chromatographic condition

Chromatographic column: Waters Acquity HSS T3 column (2.1×150 mm, 1.8 μm); Column temperature: 35°C; Mobile phase: A-water (containing 0.1% formic acid); B-acetonitrile; Flow rate: 0.3 ml/min; Injection volume: 2 μl elution gradient.

### *In vitro* cell experiment

The healthy mouse hepatocyte cell line AML-12 was obtained from Nanjing Drum Tower Hospital. AML-12 cells were cultured in Dulbecco’s modified Eagle’s medium (DMEM) (KeyGEN Biotech, Wageningen, the Netherlands) and supplemented with 1% penicillin/streptomycin and 10% fetal bovine serum (FBS; Gibco, Waltham, MA).

Ethanol injury in AML-12 cells was evaluated by assessing cell viability using the CCK-8 assay. The cells were divided into control and alcohol-treated groups. AML-12 cells (1×10^4^ cells/well) were seeded into 96-well plates and incubated for 24 h. Fresh medium and medium containing varying concentrations of ethanol (10–80 mg/mL) were added for 24 and 48 h. The ethanol concentration that resulted in 50% cell viability was then determined.

Toxicity analysis of the active ingredients of the PFL metabolites in the AML-12 cells was performed using the CCK8 assay. The experimental groups were treated with media containing varying concentrations of the active ingredients (50–2000 ng/mL).

In regards to the protective effect of 3,4-Hydroxyphenyl lactic acid on ethanol-injured AML-12 cells, cells were divided into control, model, and experimental groups. AML-12 cells (2 × 10^5^ cells/well) were incubated in a six-well plate for 24 h. The fresh medium was replaced with medium containing different concentrations of antioxidant peptides (0.1μg/mL and 0.5 μg/mL) for an additional 24 h. Subsequently, the medium containing the ethanol concentration was added, and the cells were incubated for another 24 h for follow-up experiments.

### Antioxidant activity of active components of PFL metabolites

Four types of peptides were synthesized by Tianjin Peptide Valley Biotechnology Co., Ltd. The antioxidant capacity of the metabolites was assessed using free radical scavenging assays with 2,2’-azinobis (3-ethylbenzothiazoline-6-sulfonic acid) diammonium salt and 1,1-diphenyl-2-picrylhydrazyl. Both methods were performed according to manufacturer’s instructions provided in the kit (Box Bio).

### Oil Red O Staining

AML-12 cells (1.5 × 10^5^ / well) were seeded in a six-well plate and induced by oleic acid at a final concentration of 0.2 mM for 24 h. Subsequently, the cells were fixed, rinsed, and stained with oleic acid, after which they were rinsed again and mounted for observation and photography under a 40× microscope.

### Network pharmacology

#### 1. Analysis of target genes of major metabolites in PFL

The SMILES names of peptides 74, 78, and 82, as well as HPLA, were obtained from PubChem (https://pubchem.ncbi.nlm.nih.gov/) and imported into Swiss ADME (http://www.swissadme.ch/index.php). The compounds were screened based on a gastrointestinal absorption score of ‘high’ and a druglike-ness score of at least two ‘yes’ responses. The canonical SMILES of the screened compounds were then submitted to the Swiss Target Prediction database (http://swisstargetprediction.ch/) to predict the targets of each compound. The predicted targets were further validated using the Uniprot database (https://www.uniprot.org/) to identify the potential targets of the major metabolite active ingredients in the PFL.

#### 2. Alcoholism Disease Target Analyses

Using” alcoholism’ as the keyword, we retrieved targets for alcoholism from the GeneCards (https://www.genecards.org/) and DisGeNET (https://disgenet.com/) databases. After collecting all targets, duplicate entries were removed, and the target names were normalized. A Venn diagram was generated using Venny 2.1.0 (https://bioinfogp.cnb.csic.es/tools/venny/index.html) to illustrate the intersection of the target genes between the main metabolites and alcoholism.

#### 3. Protein-protein interaction network construction

Intersection target genes were introduced into the STRING database (https://cn.string-db.org/) for protein-protein interaction analysis. The results were subsequently imported into Cytoscape 3.9.1, where the Centiscape 2.2 plug-in was used to examine the network topology. The threshold values for betweenness, closeness, and degree were selected as the core target genes, which resulted in the generation of a PPI network map.

#### 4. Gene Ontology (GO) and Kyoto Encyclopedia of Genes and Genomes (KEGG) pathway enrichment analysis

The Metascape database (https://metascape.org/) was used to analyze the enrichment of overlapping genes through GO and KEGG pathways. Using a screening threshold of p < 0.01, three modules were analyzed in the GO analysis, biological process, molecular function, and cellular component, as well as the KEGG pathways, to identify significantly enriched signaling pathways.

### Western Blot analysis

Cellular proteins were extracted and quantified. Equal amounts of proteins were separated by 10% SDS-PAGE and subsequently transferred to a polyvinylidene fluoride membrane (Bio-Rad, Hercules, CA, USA). The membrane was blocked with 5% skim milk or 5% bovine serum albumin and incubated with the primary antibody overnight. On the following day, the membrane was incubated with a secondary antibody. Chemiluminescence imaging and ECL detection reagents were used to visualize the signals. The antibodies used for western blotting are listed in Table S2.

### Statistical analysis

All results were confirmed in at least three independent experiments. Data were processed using GraphPad Prism V.9.0 software; the results are expressed as mean ± standard deviation. Statistical analyses were performed using one-way analysis of variance or Tukey’s test. Statistical significance was set at p < 0.05.

## Results

### PFL attenuates acute alcohol intoxication in mice

To investigate the direct effects of PFL on alcoholism, we developed a mouse model of acute alcoholism and analyzed the differences in physicochemical indices among three groups: pretreated with phosphate-buffered saline (Model), *L. plantarum* (P group), and PFL (PFL group) (Fig. 1A). Survival analysis revealed that the mortality rate in the PFL group was 12.5% (1/8), which was significantly lower than the 62.5% (5/8) mortality rate in the acute alcoholism model group (*P* < 0.05). In contrast, the P pretreatment group exhibited a mortality rate of 25% (2/8), which was not significantly different from that in the acute alcoholism group (Fig. 1B). Additionally, the duration of intoxication and tolerance in different groups was assessed using a flip-flop reflex experiment (Fig. 1C). The results indicated that the flip-flop reflex disappeared in the acute alcoholism model group within 15 min. However, the intoxication latency and tolerance time were significantly prolonged in mice pretreated with P and PFL (*P* < 0.001) (Fig. 1D). These findings demonstrated that both P and PFL reduced mortality rates associated with acute alcoholism, prolonged the time to alcohol tolerance, and facilitated sobriety in mice experiencing acute alcohol intoxication, with PFL exhibiting a more pronounced antidote effect compared to that of the P group.

**Figure 1.**
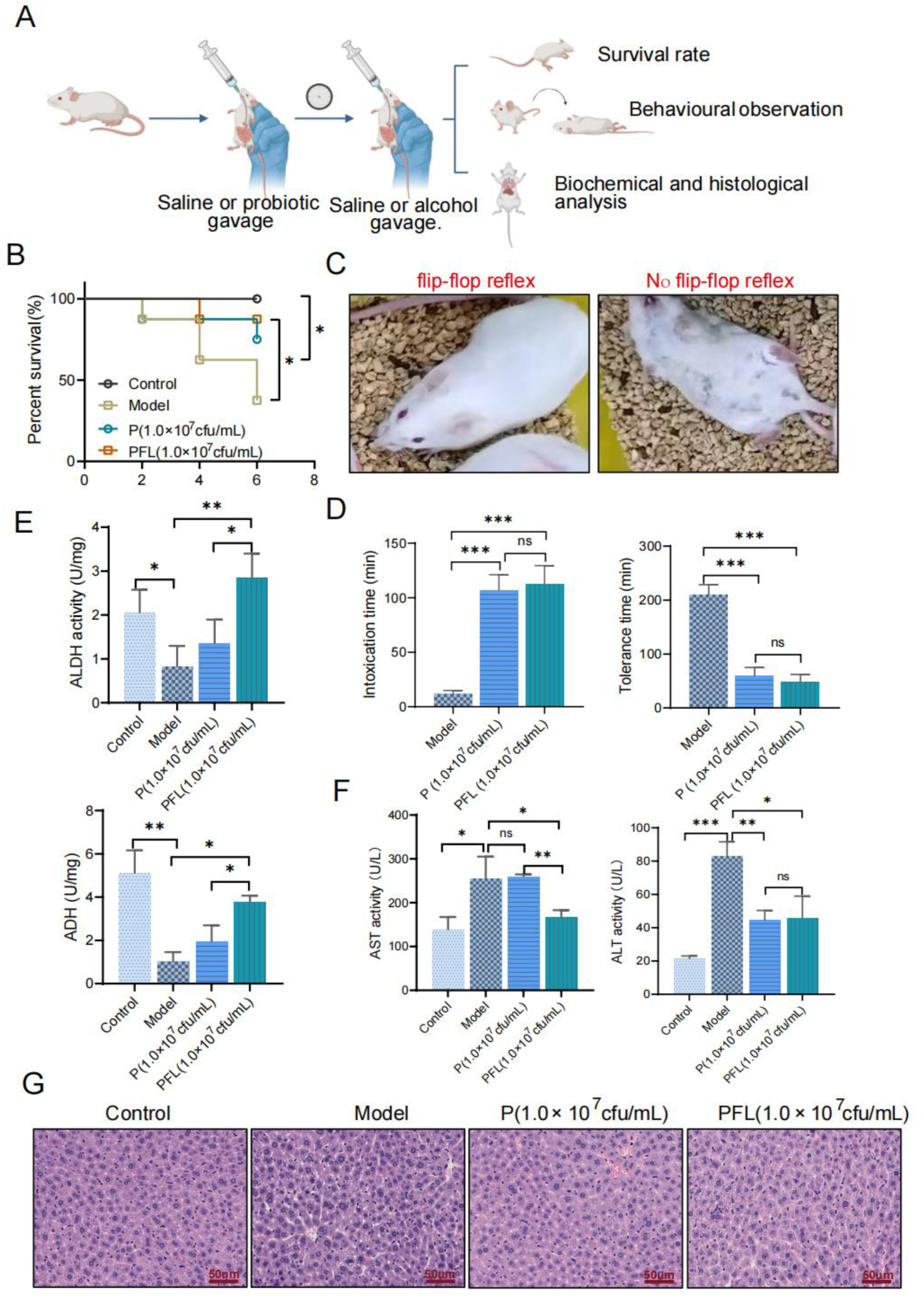
Protective effect of PFL on acute alcoholism model in mice. (A) Experimental flowchart. The mice were initially gavaged with 0.2 mL saline, P at 1×10^7^ cfu/0.2 mL, or PFL at 1×10^7^ cfu/0.2 mL. After a 30-minute interval, they received another gavage of 10 mL/kg saline or 56% ethanol. Gavage was conducted thrice every other day. Mouse survival, behavioral changes, and biochemical and histological analyses were performed 1 h after the third alcohol administration via gavage. (B) Kaplan-Meier survival curves for a mouse model of acute alcohol intoxication. (C) The general status of normal and intoxicated mice. (D) Time to onset of intoxication (duration from alcohol consumption to disappearance of the flip-flop reflex) and time to tolerance (duration from disappearance of the flip-flop reflex to the return of the flip-flop reflex). (E) Activities of ALDH (upper chart) and ADH (lower chart) in liver tissues. (F) Serum AST and ALT levels (G) H&E-stained images of typical liver tissue. *Statistical significance is indicated as follows: \**P*<0.05;\*\**P*<0.01;\*\*\**P*< 0.001; ns means *P*>0.05. Mantel-Cox log--rank test in B. Two-sided Student’s *t*-test was used to compare two groups in D, E, and F. Results are expressed as three replicate values in side-by-side subcolumns.

ALDH and ADH are the primary enzymes involved in alcohol metabolism(Liu et al. 2019). We measured the activities of ALDH and ADH in the liver tissues to investigate the effect of PFL on alcohol metabolism. The results showed a significant reduction in ALDH and ADH activity in the model group compared to that of the control group. However, the PFL pretreatment group exhibited a significant increase in ALDH and ADH activities compared to that of the model group, and the effect of the PFL pretreatment group was greater than that of the P pretreatment group (Fig. 1E). The extent of liver injury was assessed by measuring serum levels of AST and ALT. The results indicated that AST and ALT activities were significantly elevated in the model group compared with those in the control group. However, pretreatment with PFL significantly reduced AST and ALT activity in the model group (Fig. 1F). Histological analysis revealed increased inflammatory cell infiltration and a greater number of fatty vacuoles in the interstitial spaces of hepatocytes in the model group than in the control group. In contrast, hepatocyte steatosis was significantly reduced in the PFL-pretreated group (Fig. 1G), suggesting that PFL exerts a protective effect against acute alcohol-induced liver injury.

### PFL alleviates chronic alcoholic liver injury through an anti-oxidative stress mechanism

To further explore whether PFL mitigates long-term alcohol-induced injury, we established a mouse model of chronic alcohol-induced injury. The liver index, defined as the ratio of liver weight to body weight, is an important indicator of nutritional status and liver pathology in mice. Compared with the control group, the liver index was significantly higher in the model group (*P*<0.05). However, pretreatment with both P and PFL significantly reduced the liver index (*P*<0.05) (Fig. 2A). Alcohol gavage induced liver swelling in mice, whereas pretreatment with P or PFL effectively mitigated this swelling and exerted a protective effect on the liver.

**Figure 2.**
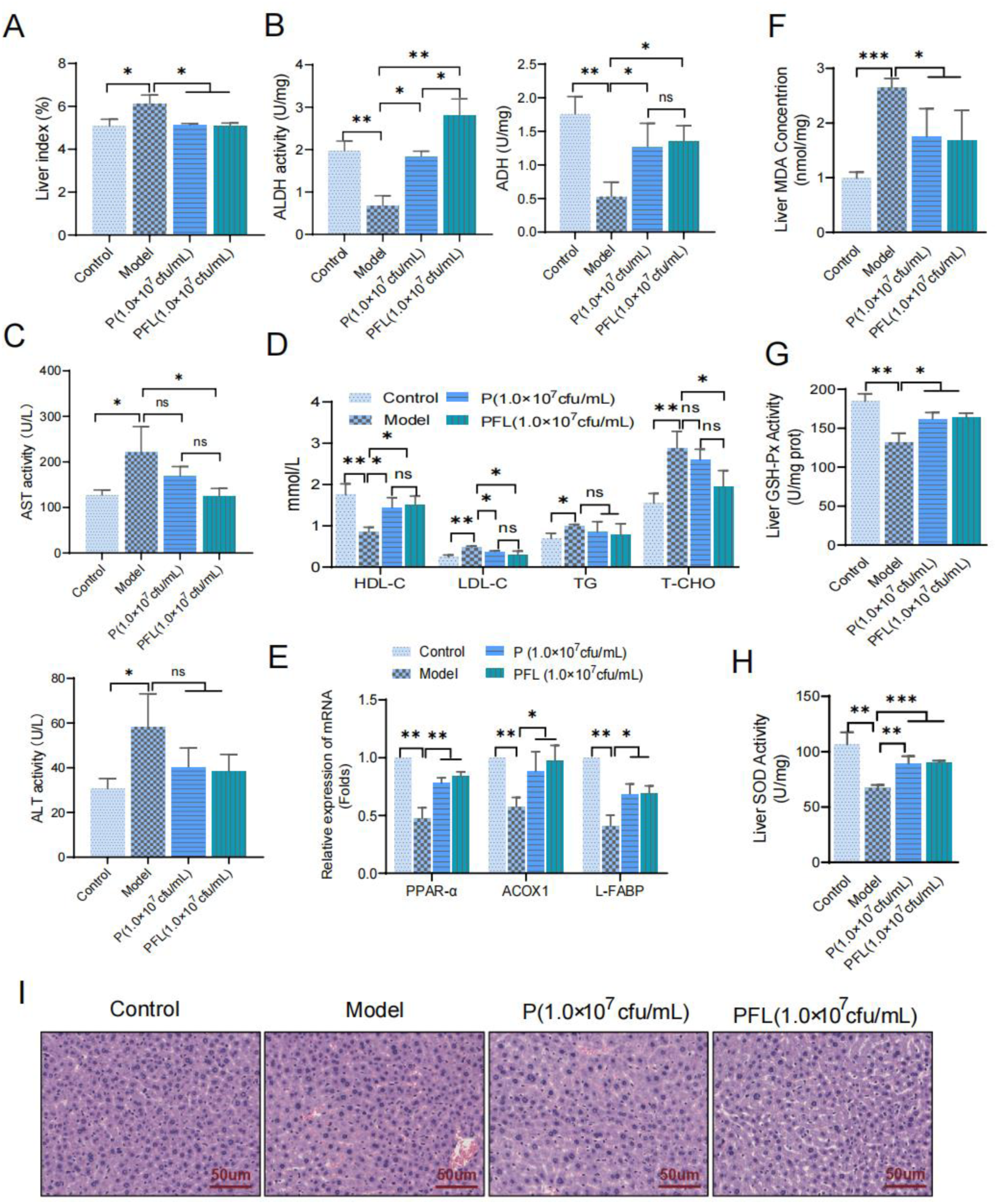
PFL alleviates chronic alcoholic liver injury through an anti-oxidative stress mechanism. The P and PFL groups were administered a gavage of 1×10^7^ cfu/0.2 mL for 30 minutes, after which the mice received a gavage of 2 mL/kg of either 56% ethanol or saline every other day for a duration of 5 weeks. (A) Liver indices of different groups of mice. (B) Activities of ALDH and ADH in liver tissues. (C) Serum activities of AST (upper chart) and ALT (lower chart). (D) Concentrations of HDL-C, LDL-C, TG, and T-CHO. (E) mRNA expression of PPAR-α pathway-related genes in mice liver tissues. (F-H) Activities of MDA, GSH-Px, and SOD in mouse liver tissue. (I) H&E-stained images of liver tissues from mice with chronic alcohol injury. *Statistical significance is indicated as follows: *P<0.05;**P<0.01;***P<0.001; ns means P>0.05. A two-sided Student’s t-test was used for comparisons between the two groups, as shown in panels A-H. The results are expressed as three replicate values in the side-by-side subcolumns.

We investigated the effects of PFL on alcohol-metabolizing enzymes and serum biochemical indices in a chronic alcohol injury model. Compared to the control group, ALDH and ADH activities were significantly decreased in the model group; however, pretreatment with P and PFL significantly elevated ALDH activity and increased ADH activity compared to the model group. The PFL-pretreated group had higher ALDH and ADH activities than the P-pretreated group (Fig. 2B). Biochemical analyses revealed that serum AST activity was enhanced in the model group compared to the control group, while AST activity was significantly decreased in mice pretreated with PFL; however, no significant difference in AST activity was observed between the P and model groups. Additionally, there was no significant difference in ALT activity between the groups (Fig. 2C). HDL-C, LDL-C, TG, and T-CHO, key indicators of lipid metabolism in fatty livers, were also measured. Compared to the control group, HDL-C levels were significantly reduced in the model group, whereas they were significantly elevated in the groups pretreated with both P and PFL; a similar trend was observed for LDL-C levels. Furthermore, TG and T-CHO levels were elevated in the model group. Although pretreatment with PFL significantly reduced the T-CHO levels, no significant differences were observed between the P and model groups (Fig. 2D). These results suggest that PFL pretreatment exerts a more pronounced protective effect against alcoholic liver injury than pretreatment with P, and that it does this by balancing lipid levels and reducing liver damage.

Oxidative stress plays a critical role in the progression of alcoholic liver injury and hepatocyte damage is closely linked to free radical production. Peroxisome proliferator-activated receptor alpha (PPAR-α), a ligand-activated transcription factor, belongs to the nuclear hormone receptor (NR) superfamily and is associated with fatty acid transport, mitochondrial fatty acid oxidation, inflammatory responses, and fibrogenesis. PPAR-α is a key regulator of hepatic oxidative stress. Acyl-CoA oxidase 1 (ACOX1) and liver fatty acid-binding protein (L-FABP) are downstream target genes of PPAR-α. ACOX1 promotes fatty acid metabolism, whereas l-FABP regulates fatty acid uptake and intracellular transport.

We extracted RNA from mice liver tissues, and QPCR analysis revealed that the mRNA levels of PPAR-α, ACOX1, and L-FABP were significantly reduced in the liver tissues of the model group compared to control mice. In contrast, pretreatment with P and PFL significantly increased the mRNA levels of PPAR-α, ACOX1, and L-FABP, with PFL showing a more pronounced effect than P (Fig. 2E). These findings suggest that PFL may mitigate lipid metabolism disorders and alleviate alcoholic liver injury by selectively regulating the PPAR-α pathway and its associated genes.

To further investigate the effect of PFL on the antioxidant capacity in the liver of mice with chronic alcoholism, we measured the levels of MDA, GSH-Px, and SOD. The result showed that MDA activity in the model group was significantly higher than in the control group (*P*<0.01). In contrast, GSH-Px and SOD activities were significantly lower in the model group than in the control group (*P*<0.001). Pretreated with both P and PFL resulted in significantly lower MDA levels compared to the model group (*P*<0.05) (Fig. 2F), whereas GSH-Px and SOD activities were significantly higher than those in the model group (Fig. 2G-H). These findings indicate that PFL mitigates hepatocyte damage caused by alcohol-induced oxidative stress from alcohol exposure by enhancing antioxidant enzyme activity and reducing lipid peroxidation.

Histological analysis revealed that, compared to the control group, hepatocytes in the model group had an irregular morphology with disorganized hepatic cords and a large accumulation of fat vacuoles. In contrast, the hepatocyte morphology in the PFL pretreatment group was more regular, with reduced steatosis (Fig. 2I). These findings suggest that PFL exerts protective effects against chronic alcoholic liver injury.

### Identification of PFL components through UHPLC-QTOF-MS/MS

To further analyze the components of PFL, UHPLC-Q-TOF-MS/MS was used to identify a wide range of amino acids, dipeptides, and peptides, nucleosides such as guanine, guanine nucleotides, cyclic guanosine monophosphate, and other compounds such as vitamin B6 (pyridoxamine), fatty acids, and lactic acid (Table S3, Fig. 3A-B). Four substances with high expression levels, polypeptides 74, 78, and 82, and HPLA, were selected for structural analysis and synthesis (Fig. 3C), and their functions were further verified.

**Figure 3.**
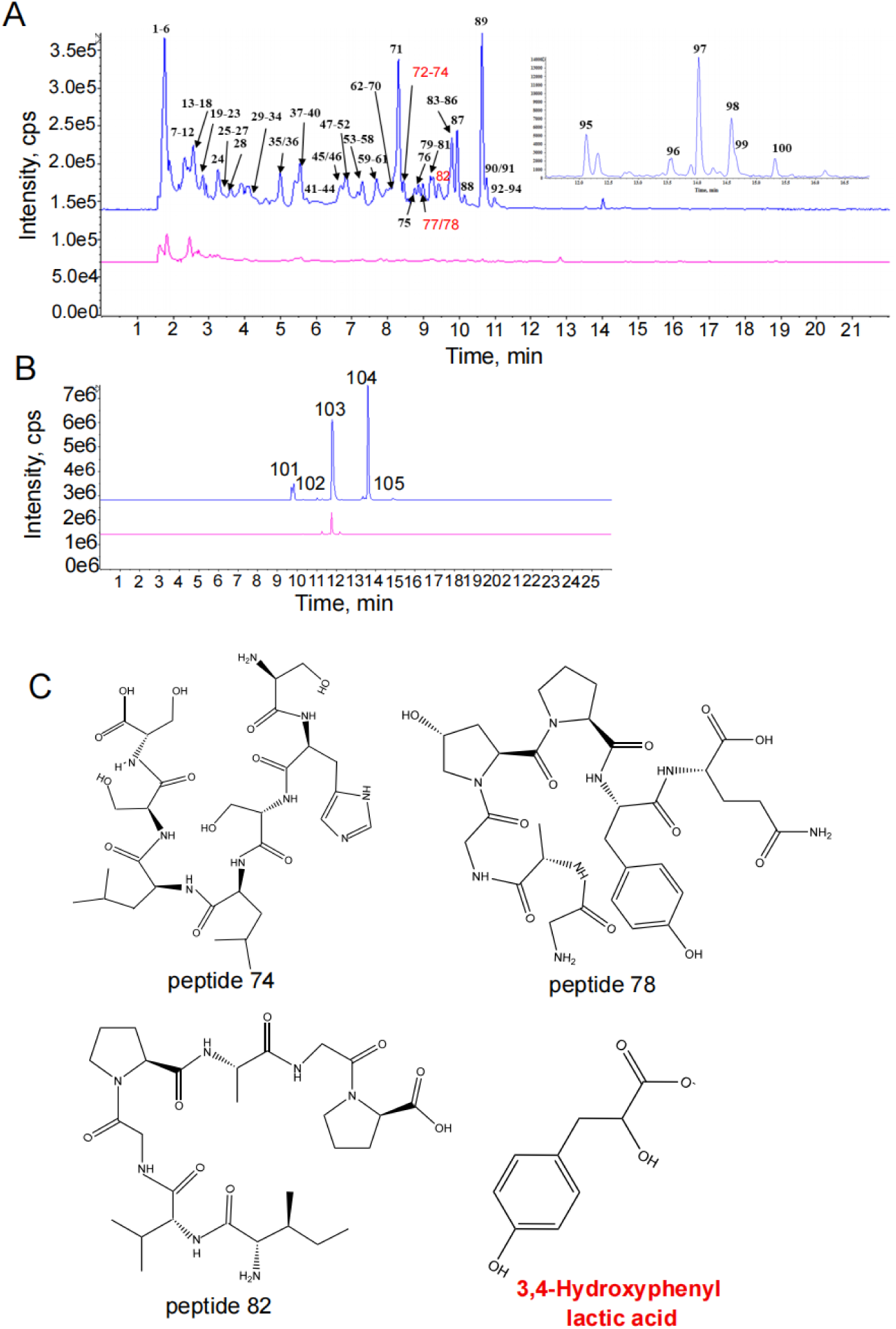
Identification of PFL components through ultra-high performance liquid chromatography-quadrupole-time of flight mass spectrometry (UHPLC-QTOF-MS/MS) (A) The underived combined extracted ion flow diagrams (with positive and negative ions); blue represents the post-fermentation sample and red represents the pre-fermentation sample. (B) The derivatization combined extracted ion flow patterns (with positive and negative ions); blue represents the post-fermentation sample and red represents the pre-fermentation sample. (C) Structures of the four active peptides.

### HPLA alleviates alcoholic liver injury through antioxidant stress

Ethanol induces oxidative stress, diminishes antioxidant capacity, and contributes to alcoholism. Therefore, compounds exhibiting antioxidant activity may offer protection against ethanol-induced alcoholic liver injury. We assessed the antioxidant activity of the four active ingredients and found that polypeptide 78 and HPLA demonstrated significant antioxidant effects with a concentration-dependent enhancement trend. Among these, HPLA exhibited the highest antioxidant capacity (Fig. 4A-B). Subsequently, we treated the normal mouse liver cell line (AML-12) with varying concentrations of peptide 78 (50-2000 ng/mL) and HPLA (50-2000 ng/mL) for 24 and 48 h. Network pharmacological analysis revealed that peptide 78 lacked drug-like properties, whereas HPLA exhibited favorable drug-like characteristics (Figure S1A-B). Therefore, we selected HPLA for further analysis.

**Figure 4.**
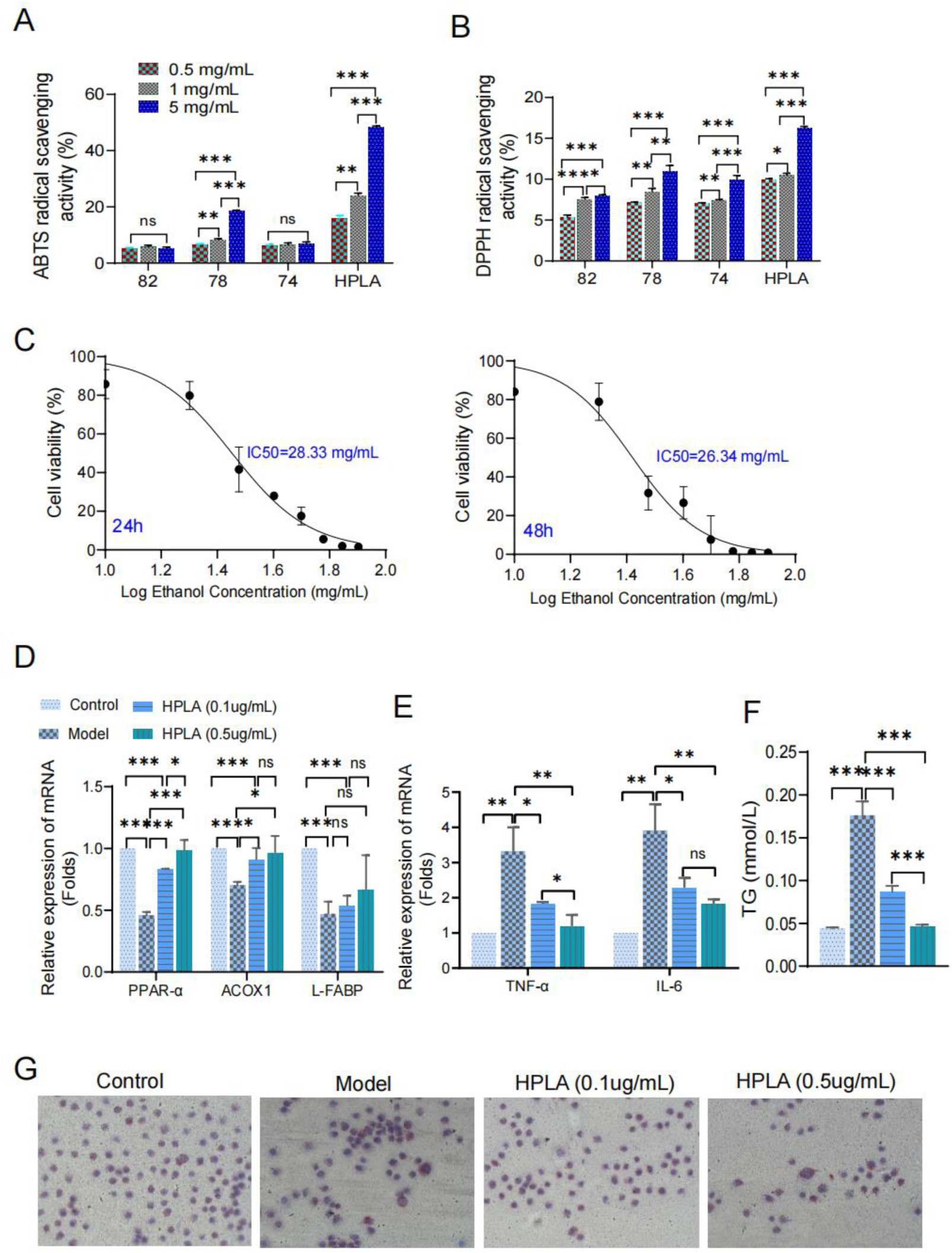
HPLA alleviates alcoholic liver injury through antioxidant stress. (A–B) Antioxidant activity of active components from PFL metabolites assessed using the ABTS and DPPH methods. (C) Cell viability of AML-12 cells after treatment with different concentrations of alcohol for 24 and 48 hours. (D) Expression of mRNA related to the PPAR-α pathway in cells. (E) Expression of tumor necrosis factor-alpha (TNF-α) and interleukin 6 (IL-6) mRNA in cells. (F) Determination of triglyceride (TG) content in cells. (G) Image of oil red O staining results in AML-12 cells. *Statistical significance is indicated as follows: \**P*<0.05;\*\**P*<0.01;\*\*\**P*< 0.001; ns means *P*>0.05. The two-sided Student’s *t*-test was used for comparisons in A–B and D–F; for C, values were transformed using [X = Log(X)]. Results are expressed as three replicate values in side-by-side subcolumns.

AML-12 cell viability decreased as the ethanol concentration increased in the ethanol injury model construct. The ethanol concentration reached approximately 30 mg/mL when the cell survival rate was 50%, 24 and 48 h after ethanol induction (Fig. 4C). Therefore, we constructed an alcohol damage model in mouse liver cells using 30 mg/mL ethanol to evaluate the protective effects of HPLA against oxidative damage in AML-12 cells. In this study, the mRNA levels of PPAR-α, ACOX1, and L-FABP in the model group were significantly lower than those in the control group. In comparison to the model group, the mRNA levels of PPAR-α and ACOX1 in the HPLA group were significantly higher; however, there was no significant difference in L-FABP levels (Fig. 4D). The results suggested that HPLA may alleviate alcoholic liver injury by regulating the expression of the PPAR-α pathway and its related genes, with a concentration of 0.5 μg/mL proving to be the most effective. Furthermore, the mRNA levels of TNF-a and IL-6 were significantly higher in the model group compared to the control group, whereas pretreatment with HPLA significantly reduced the mRNA levels of these inflammatory factors (Fig. 4E). The accumulation of TG in the liver is a critical factor in the progression from simple steatosis to steatohepatitis. In the model group, the intracellular TG content was significantly higher than that in the control group (*P*<0.001). Pretreatment with two different doses of HPLA significantly reduced the TG content in the model group (*P* < 0.001), with the 0.5 µg dose demonstrating a more pronounced effect compared to the 0.1 µg dose (*P*<0.001) (Fig. 4F). Consistently, the results of Oil Red O staining revealed that a substantial number of red lipid droplets were formed in the cells of the model group, arranged in a continuous bead-like pattern with clear boundaries. In contrast, HPLA treatment led to a noticeable reduction in the intracellular lipid droplets, which appeared to be unevenly distributed and arranged in a disordered manner (Fig. 4G). These findings indicated that HPLA can effectively reduce lipid droplet formation and alleviate alcohol-induced fatty liver lesions in vitro.

### HPLA alleviates alcoholic liver injury by interfering with the EGFR/PPAR-α oxidative stress signaling axis

First, we used the Swiss Target Prediction database to identify a total of 100 drug targets and 16 potential drug targets for HPLA. Subsequently, we gathered 1,736 disease targets related to alcoholism using the GeneCards and DisGeNET databases. Drug and disease targets were analyzed using a Venn diagram, which revealed nine intersecting targets (Fig. 5A). These nine targets were then imported into the STRING database, resulting in a protein-protein interaction (PPI) network graph comprising six nodes and ten edges (Fig. 5B). This PPI network was subsequently imported into Cytoscape 3.9.1 for network topology analysis, using the Centiscape 2.2 plug-in. Ultimately, the core targets identified were ESR1, PPARG, PPARA, EGFR, FYN, and PPARD (Fig. 5C).

**Figure 5.**
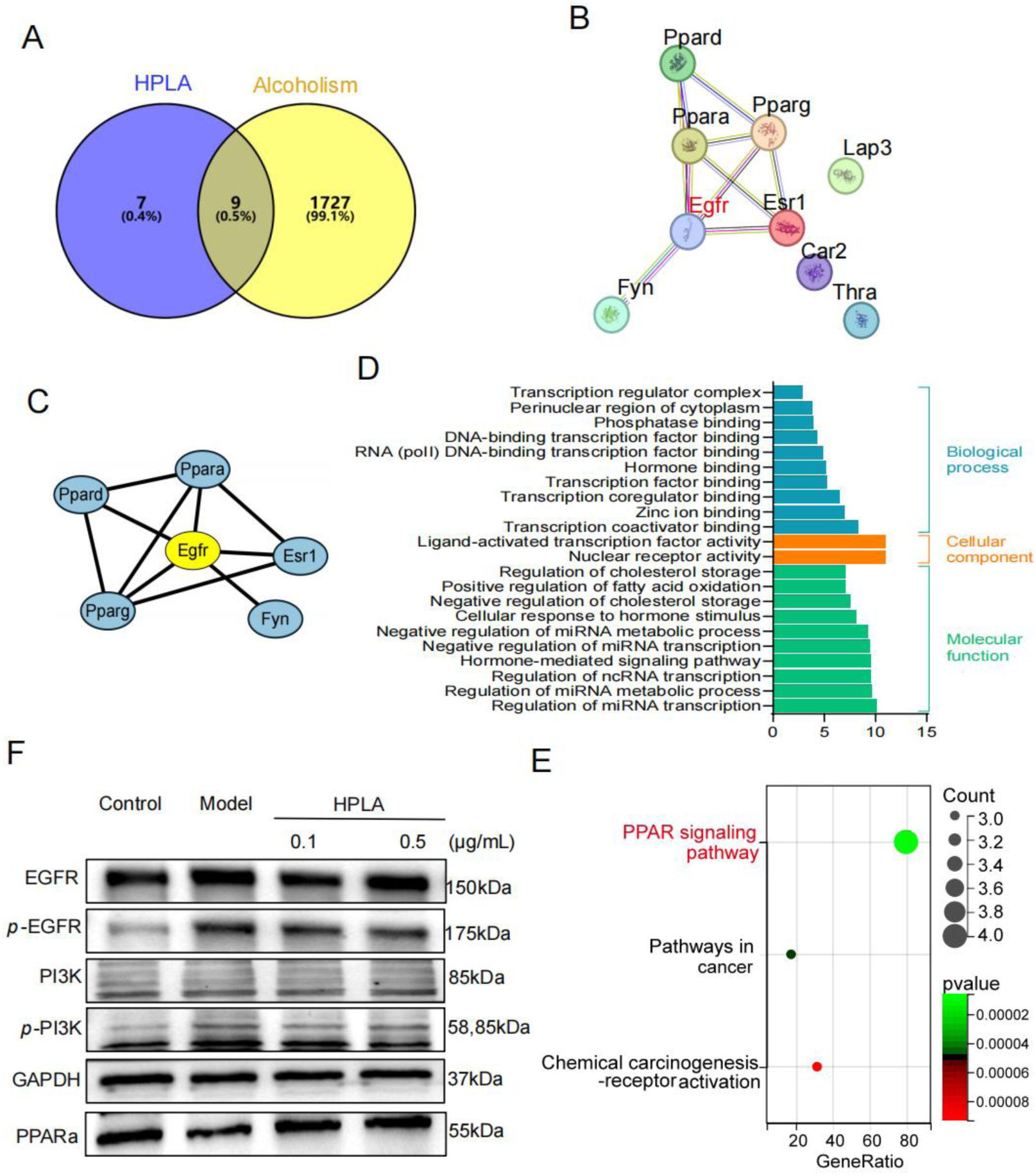
HPLA alleviates alcoholic liver injury by interfering with the EGFR/PPAR-a oxidative stress signaling axis. (A) Venn diagram illustrating the intersection of HPLA targets with those associated with alcoholism. (B) Protein-protein interaction network diagrams of intersecting targets. (C) Topological analysis of core intersection targets. (D) Triple histograms displaying GO term enrichment of intersecting targets in biological processes, cellular components, and molecular functions. (E) Bubble diagram representing KEGG pathway enrichment analysis of intersecting targets. (F) Western blot analysis showing the levels of phosphorylated EGFR and PI3K (*p*-EGFR and *p*-PI3K) as well as total EGFR and PI3K in the AML-12 cell model of alcoholism.

In addition, we conducted GO and KEGG enrichment analyses of the nine core targets of HPLA for the treatment of alcoholism. GO enrichment analysis revealed 104 enrichment results associated with biological processes, 22 with cellular components, and 16 with molecular functions. Among these findings, we filtered the top 10 results (Fig. 5D). The biological processes were primarily related to the negative regulation of miRNA transcription, metabolic processes, ncRNA transcription, hormone-mediated signaling pathways, cellular responses to hormonal stimuli, negative regulation of cholesterol storage, and positive regulation of fatty acid oxidation. The cellular components mainly comprised cytoplasmic perinuclear regions and transcriptional regulatory complexes. The molecular functions involved nuclear receptors, ligand-activated transcription factor activity, transcriptional coactivators, zinc ion binding, transcriptional coregulators, transcription factors, RNA polymerase II (RNA Pol II), DNA-binding transcription factors, DNA-binding transcription factors, and phosphatase binding. KEGG enrichment analysis revealed three signaling pathways associated with oxidative damage (Fig. 5E), primarily the PPAR signaling pathway, pathways in cancer, and the chemical carcinogenesis-receptor activation pathway. These findings suggest that HPLA exerts its antioxidant effects by influencing both cancer-related and PPAR signaling pathways. Furthermore, HPLA appears to attenuate alcohol-induced oxidative stress damage by regulating biological processes across multiple cellular compartments and functions.

Previous web-based pharmacological analyses indicated that EGFR is a key target of HPLA for the treatment of alcoholism, with PI3K serving as the major downstream signaling molecule. In this study, Western blot analysis revealed that alcohol activated the EGFR and PI3K signaling pathways, whereas treatment with HPLA significantly inhibited their activation. Conversely, alcohol inhibited the activation of PPAR-α, but HPLA treatment mitigated the inhibitory effects of alcohol on PPAR-α (Fig. 5F). These results align with previous findings from the in vivo mice model, suggesting that HPLA may alleviate alcohol-induced oxidative stress injury by modulating the EGFR/PPAR-α pathway.

### HPLA alleviates acute alcoholic liver injury in mice

To further verify the protective effect of HPLA in the alcohol intoxication model, we re-established an acute alcohol intoxication mouse model by intravenous injection (Fig. 6A). Compared to the model group, pretreatment with HPLA significantly prolonged the time to develop alcohol intoxication and reduced the duration of intoxication in mice (Fig. 6B). Analysis of alcohol metabolism enzymes in vivo showed that the activities of ALDH and ADH in the liver tissues of the model group were significantly reduced compared to those in the control group. However, the alcohol-induced decreases in ALDH and ADH activities were partially restored after pretreatment with HPLA (Fig. 6C). Serum biochemical analysis revealed that AST activity was significantly increased in the model group compared to that in the control group, while pretreatment with HPLA partially reduced the elevated AST activity in the model group; however, no statistical significance was found in ALT activity detection (Fig. 6D). In conclusion, these results suggested that HPLA exerts protective effects against acute alcoholism.

**Figure 6.**
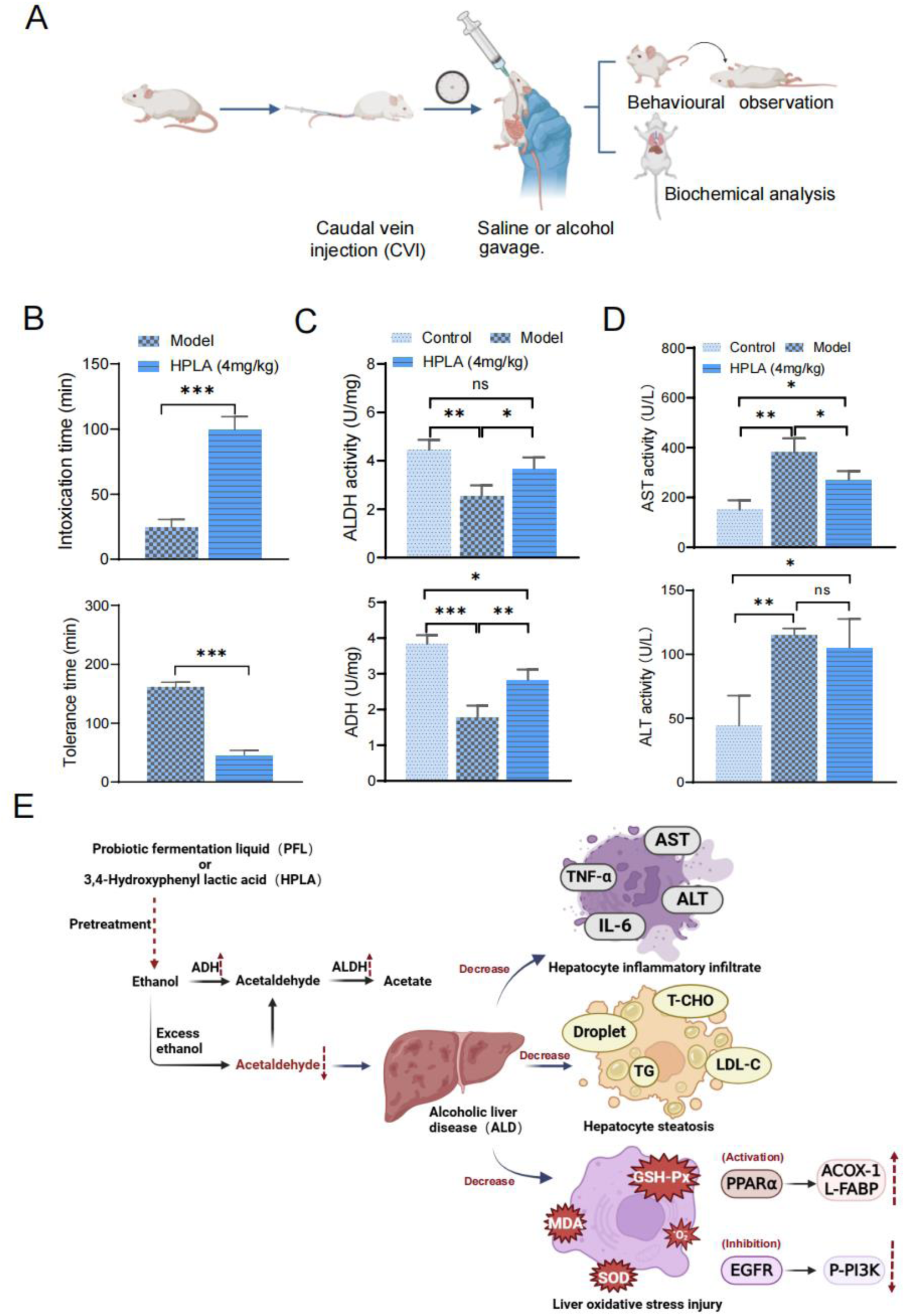
HPLA alleviates acute alcoholic liver injury in mice. (A) Experimental Flowchart: Mice were intravenously injected with 4 mg/kg HPLA or an equivalent volume of saline. Thirty minutes later, the mice were administered 10 mL/kg saline or 56% ethanol by gavage. This gavage was performed every other day for a total of three doses. Behavioral observations and biochemical analyses of liver tissues were conducted separately. (B) Intoxication and tolerance times in mice. (C) Activity of ALDH and ADH in the liver tissue. (D) Serum activity of AST and ALT levels. *Statistical significance is indicated as follows: \**P*<0.05;**P<0.01;\*\*\**P*<0.001; ns means *P*>0.05. A two-sided Student’s *t*-test was used for comparisons between the two groups (panels B–D). The results are expressed as three replicate values in the side-by-side subcolumns. (E) Schematic representation of the study.

## Discussion

Alcoholic liver injury, stemming from chronic excessive alcohol consumption, can progress through stages of fatty liver disease, hepatitis, fibrosis, and cirrhosis, underscoring the need for effective therapeutic interventions (Liu et al. 2019). *Lactiplantibacillus plantarum* (*L. plantarum*), a probiotic with demonstrated benefits in modulating gut microbiota, enhancing immunity, and improving metabolic health (Chahwan et al. 2019; Kim et al. 2019; Alli et al. 2022), has emerged as a promising candidate for mitigating alcohol-induced liver damage. Notably, Arora et al. (2014) highlighted the synergistic effects of Bifidobacterium and *L. plantarum* in restoring gut-liver axis homeostasis and reducing hepatic inflammation via IL-12/p40 inhibition. However, the specific molecular mechanisms underlying *L. plantarum*-mediated hepatoprotection remain incompletely elucidated.

In this study, we demonstrated that *L. plantarum* fermentation liquid (PFL) conferred robust protection against both acute and chronic alcohol-induced liver injury in murine models, outperforming *L. plantarum* alone. PFL, enriched with metabolites such as organic acids, antimicrobial agents, and short-chain fatty acids (Xiao et al. 2022; Li et al. 2023), likely exerts its benefits through a combination of bioactive compounds. UHPLC-QTOF-MS/MS analysis identified 3,4-hydroxyphenyl lactic acid (HPLA) as a principal active component in PFL, exhibiting superior antioxidant capacity compared to co-detected peptides. This aligns with prior reports of HPLA’s broad biological activities, including anti-inflammatory and antimicrobial effects (Ernst et al. 2022; Mu et al. 2010; Hughes et al. 2022). Our findings further revealed that HPLA attenuated alcohol-induced oxidative stress by enhancing hepatic antioxidant enzyme activity (SOD, GSH-Px), reducing lipid peroxidation (MDA), and mitigating lipid accumulation in hepatocytes. Despite its therapeutic promise, further investigations into HPLA’s pharmacokinetics and long-term safety are warranted to advance clinical translation.

Mechanistically, network pharmacology and experimental validation implicated the EGFR/PPAR-α signaling axis as central to HPLA’s protective effects. Alcohol exposure activated EGFR/PI3K phosphorylation while suppressing PPAR-α, exacerbating oxidative stress and lipid dysregulation. HPLA pretreatment reversed these effects, inhibiting EGFR/PI3K activation and restoring PPAR-α expression. PPAR-α, a critical regulator of fatty acid oxidation and lipid homeostasis (Zhao et al. 2019), synergizes with EGFR modulation to counteract alcohol-induced hepatic damage. These findings are consistent with prior work showing *L. plantarum*-mediated neuroprotection via EGFR pathway regulation (Shukla et al. 2020), yet our study uniquely identifies HPLA as the key effector within the gut-liver axis.

In conclusion, this work establishes HPLA, a metabolite of *L. plantarum*, as a novel therapeutic agent against alcoholic liver injury. By targeting the EGFR/PPAR-α oxidative stress pathway, HPLA alleviates lipid metabolism disturbances and hepatocyte damage, providing mechanistic insights for probiotic-based interventions. These findings not only advance our understanding of *L. plantarum*’s hepatoprotective role but also highlight HPLA’s potential as a targeted therapy for alcoholic liver disease. Future studies should focus on optimizing HPLA delivery and validating its efficacy in clinical settings.

## Acknowledgements

We thank Nanjing First Hospital for providing patient tumor samples and technical support.

## Authorship contribution

C Z: Writing – original draft, contributed to the data analysis, contributed to the design of animal experiments, performed the majority of the experiments.Y C: Participated in sample collection.S H: Formal analysis. X B: Formal analysis. B Y: Investigation. S L: Investigation. X F: Investigation. S Z, Y P and W Z: Writing – review & editing, Funding acquisition, Conceptualization.

## Supplementary material

Supplementary data are available at *Applied Microbiology and Biotechnology* online.

## Availability of data and materials

The data obtained and/or analyzed during the current study were available from the corresponding authors in a reasonable request.

## Declarations

### Ethics approval

All animals were handled in accordance with the Guide for the Care and Use of Laboratory Animals (NIH Publication No. 85-23, Revised 1996). Animal experiments were approved by the Animal Care Committee of the Nanjing First Hospital, Nanjing Medical University.

